# THE YKI-CACTUS (I_K_Bα)-JNK AXIS PROMOTES TUMOR GROWTH AND PROGRESSION IN DROSOPHILA

**DOI:** 10.1101/2020.08.16.253005

**Authors:** Kirti Snigdha, Amit Singh, Madhuri Kango-Singh

**Affiliations:** Department of Biology, University of Dayton, Dayton OH 45469; Center for Tissue Regeneration and Engineering at Dayton (TREND), University of Dayton, Dayton OH 45469; Premedical Programs, University of Dayton, Dayton OH 45469; Integrative Science and Engineering Center (ISE), University of Dayton, Dayton OH 45469

**Keywords:** Cancer, Yki, Ras, Inflammation, Signaling pathway, Proliferation, Invasion

## Abstract

Presence of inflammatory factors in the tumor microenvironment is well known yet their specific role in tumorigenesis is elusive. The core inflammatory pathways are conserved in *Drosophila*, including the Toll-Like Receptor (TLR) and the Tumor Necrosis Factor (TNF) pathway. We used *Drosophila* tumor models to study the role of inflammatory factors in tumorigenesis. Specifically, we co-activated oncogenic forms of *Ras^V12^* or its major effector Yorkie (*Yki^3SA^*) in polarity deficient cells mutant for tumor suppressor gene *scribble* (*scrib*) marked by GFP under *nubGAL4* or in somatic clones. This system recapitulates the clonal origins of cancer, and shows neoplastic growth, invasion and lethality. We investigated if TLR and TNF pathway affect growth of *Yki^3SA^scrib^RNAi^* or *Ras^V12^scrib^RNAi^* tumors through activation of tumor promoting Jun N-terminal Kinase (JNK) pathway and its target Matrix Metalloprotease1 (MMP1). We report, TLR component, Cactus (Cact) is highly upregulated in *Yki^3SA^scrib^RNAi^* or *Ras^V12^scrib^RNAi^* tumors. *Drosophila* Cactus (mammalian I_K_Bα) acts as an inhibitor of NF_K_B signaling that plays key roles in inflammatory and immune response. Here we show an alternative role for Cactus, and by extension cytokine mediated signaling, in tumorigenesis. Downregulating Cact affects both tumor progression and invasion. Interestingly, downregulating TNF receptors in tumor cells did not affect their invasiveness despite reducing JNK activity. Genetic analysis suggested that Cact and JNK are key regulators of tumor progression. Overall, we show that Yki plays a critical role in tumorigenesis by controlling Cact, which in turn, mediates tumor promoting JNK oncogenic signaling in tumor cells.

## Introduction

Oncogenic forms of Ras are dominant drivers of tumor growth in a third of human cancers (1). Oncogenic Ras activates multiple downstream pathways like PI3K, RAF-MEK-MAPK, JNK, p38 MAPK during tumorigenesis (2). *Drosophila* tumor models of oncogenic Ras (*Ras^V12^*) with *scribble (scrib*) – the apical basal polarity regulator (referred as *Ras^V12^scrib*^-^ tumors), show neoplastic growth, invasion and lethality (3, 4). In *Ras^V12^ scrib^-^* tumors, Eiger (Egr), the ligand of *Drosophila* TNF, induces Jun N-terminal Kinase (JNK) pathway to promote tumor cell proliferation (5, 6). In contrast, Egr mediates elimination of defective cells through JNK specifically in *scrib* mosaic clones (7). Inflammatory components from the Toll Like Receptor (TLR) and Tumor Necrosis Factor (TNF) pathways play a significant role in tumor progression through poorly understood mechanisms (8, 9). However, the complex signaling initiated by oncogenic Ras presents challenges to deciphering the role of key inflammatory signals in tumor growth and progression.

A key effector of oncogenic Ras signaling is the Hippo pathway transcriptional coactivator protein Yki (*Drosophila* YAP ortholog) (10–12). Furthermore, elevated YAP activity is linked to neoplastic behavior in cancer cells, and poor prognosis in several cancers (13–15). To study the role of these inflammatory pathways in tumorigenesis, we established a Yki dependent model by expressing activated form of Yki (*UASYki^3SA^*) in cells where *scrib* is downregulated (*UASscrib^RNAi^*) using the GAL4-UAS system or in ‘flp-out’ clones. *Yki^3SA^scrib^RNAi^* tumors show key hallmarks of cancer such as sustained proliferation via JNK activation (pJNK), Matrix Metalloprotease 1 (MMP1) mediated invasion, and degradation of basement membrane due to loss of Laminin. Using these models, we show that the *Drosophila* I_K_Bα orthologue Cactus (Cact), a member of the TLR pathway (16), is upregulated in both *Yki^3SA^scrib^RNAi^* and *Ras^V12^scrib^RNAi^* clones. Downregulation of Cact (*UAS-cact^RNAi^*) in the tumor clones results in downregulation/suppression of pJNK and MMP1. We show that Cact acts upstream of JNK, and JNK activation may occur independent of Wengen (Wgn) and Grindelwald (Grnd), the *Drosophila* TNF Receptors that signal through the JNK pathway (17). Furthermore, we report a novel Yki-Cact-JNK signaling axis that promotes tumorigenesis downstream of Yki activation that is critical in promoting Ras or Yki mediated tumor growth. The Yki-Cact-JNK axis plays a critical role, as downregulation of Yki or Sd (*Drosophila* TEAD family transcription factor), or Cact or JNK disrupts this signaling axis and inhibits *Ras^V12^scrib^RNAi^* tumorigenesis.

## Results

### Comparison of oncogenic Yki and Ras dependent tumor models

We coexpressed oncogenic Yki (*UASYki^3SA^*), *scrib^RNAi^* (*UASscrib^RNAi^*) and GFP (*UASGFP*) using *nub-Gal4* in wing imaginal discs [*nub>GFPYki^3SA^scrib^RNAi^*] (**Fig. 1A**)(18). Compared to wild-type [*nub>GFP*], the *Yki^3SA^scrib^RNAi^* larvae appeared bloated, with overgrown wing discs (**Fig. 1A’**), entered an extended larval life, and were 100% pupal lethal. The dissected wing discs revealed large neoplastic growths confined to the wing pouch (detected by GFP expression) (**Fig. S1A**). These neoplasms were invasive (**Fig. 1B**), as tumor cells (GFP positive) extruded to the basal side and breached the basement membrane labelled with Laminin (red) (**Fig. 1B, B’**)(3, 19). The *nub>Yki^3SA^scrib^RNAi^* system produced massive multilayered tumors with high lethality, therefore, in parallel we made heat-shock mediated ‘Flp-out’ clones driven by *Act>y+>Gal4* (20, 21). The wild-type clones detected by co-expression of GFP (*AyGal4>UASGFP*) in the epithelial monolayer showed jagged clone boundaries (**Fig. S1B, C**)(22), whereas the *Yki^3SA^scrib^RNAi^* clones (*AyGal4>UASGFP,UASYki^3SA^,UASscrib^RNAi^*) presented as large epithelial outgrowths with well-sorted smooth borders (**Fig. 1C**). Compared to wild-type clones (thickness 11μm), the *Yki^3SA^scrib^RNAi^* clones are multilayered (23.17μm) (**Fig. S1C**). Under comparable experimental conditions, the *Ras^V12^scrib^RNAi^* larvae (120h AEL) looked thin and sluggish, and dissected imaginal discs were fragile, neoplastic, and comprised exclusively of GFP-positive cells. When the heat shock was reduced to 3min, *Ras^V12^scrib^RNAi^* ‘flp-out’ clones also showed multi-layered organization (**Fig. S1C**). JNK mediated pro-proliferative signaling promotes tumorigenesis (8, 23–25). Consistent with this, the JNK reporter *puc^E69^*-lacZ (26) was induced in *Ras^V12^scrib^RNAi^* and *Yki^3SA^scrib^RNAi^* clones (**Fig. 1C, D**). Furthermore, pJNK levels are also upregulated in *Ras^V12^scrib^RNAi^* and *Yki^3SA^scrib^RNAi^* clones (**Fig. S1D, E**). Next, to assess invasive capacity we assessed MMP1expression in our tumor models (27–30). MMP1 expression is very low in wild-type clones (**Fig S1F**), however, MMP1 is robustly induced in both *Yki^3SA^scrib^RNAi^* and *Ras^V12^scrib^RNAi^* tumor clones (**Fig. 1E, F**). Quantification of the integrated pixel intensity of MMP1 expression in *Yki^3SA^scrib^RNAi^* clone showed a 4-fold increase compared to the surrounding normal cells (**Fig. 1G**). Taken together, these data suggest that *nub>Yki^3SA^scrib^RNAi^* and *Ay>Yki^3SA^scrib^RNAi^* clones recapitulate key tumorigenic features of the well-established *Ras^V12^scrib^RNAi^* model like increased proliferation, formation of multilayered invasive neoplasms capable of basement membrane degradation, and induction of JNK signaling. Using these models, we investigated the role of key inflammatory pathways (TLR and TNF) in tumorigenesis.

**Figure 1:**
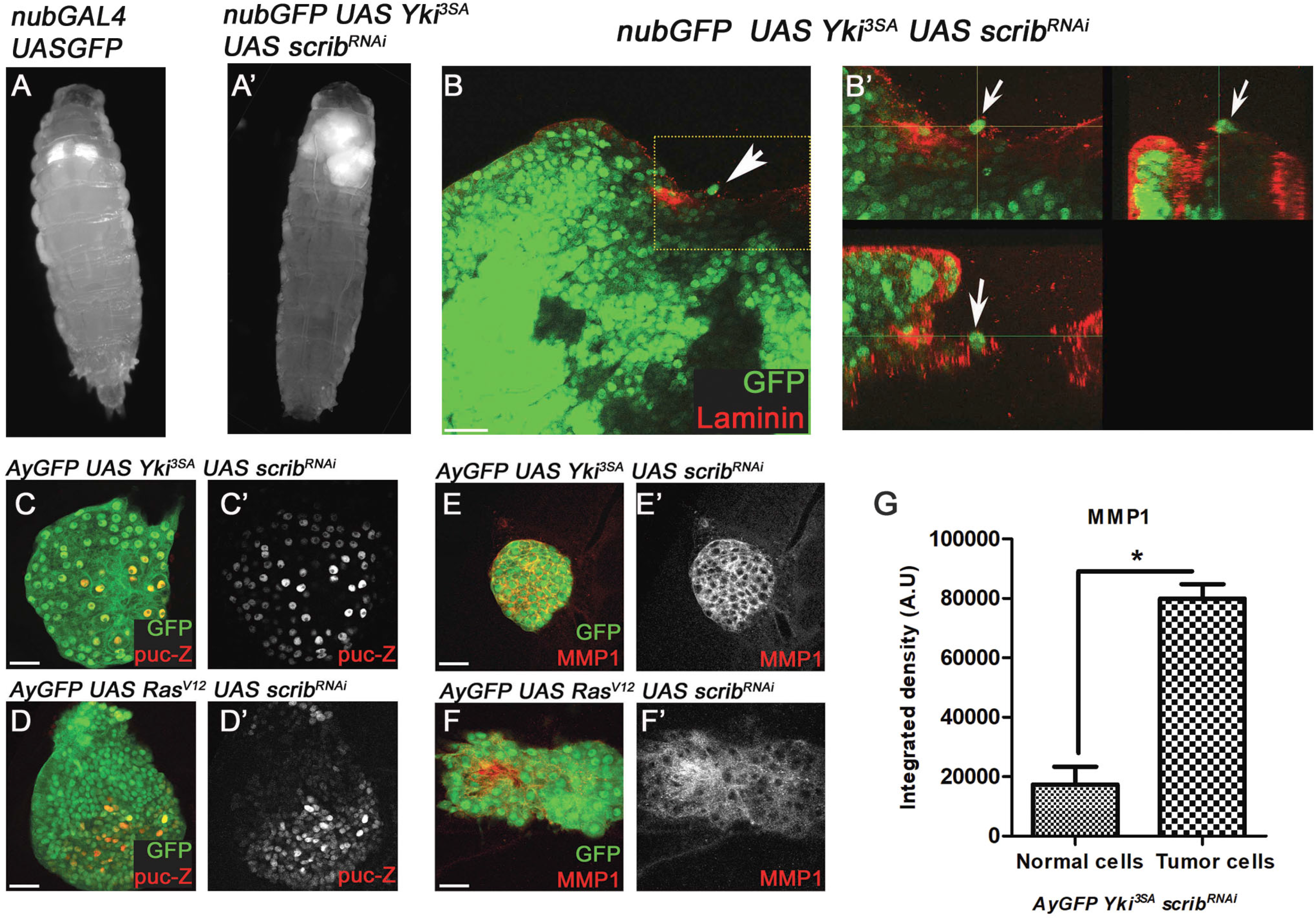
Expression of Oncogenic Yki (*Yki^3SA^*) in *scrib* mutant cells forms invasive tumors. (**A-A’**) Mature 3^rd^ instar larvae from *nub>GFP* (**A**) and *nub>Yki^3SA^scrib^RNAi^* (**A’**) show growth of wing discs (GFP, grey). (**B-B”**) Panels show (**B**) rabbit anti-Laminin (red) expression in *nub>Yki^3SA^scrib^RNAi^* (GFP, green) wing disc (60X magnification). The inset (**B’**) shows discs labelled by Laminin (red) to test motility of tumor cells (green). Panel (**B’’**) shows the XZ and YZ sections of the panel (**B**) highlighting the invasive cell (arrow). (**C-F**) Expression of *puc^E69^-lacz* tracked with mouse anti-β-gal antibody (red, grey) in (**C**) *Yki^3SA^scrib^RNAi^puc^E69^* and (**D**) *Ras^V12^scrib^RNAi^puc^E69^* clones (green). Panels show expression of MMP1 labelled with mouse anti-MMP1 antibody (red, grey) in (**E**) *Yki^3SA^scrib^RNAi^* and (**F**) *Ras^V12^scrib^RNAi^* clones (green). (**G**) Graph shows quantification of MMP1 expression in *Yki^3SA^scrib^RNAi^* tumor (GFP expressing clone) and normal cells (non-GFP expressing cells outside the clone). Paired student T test with n=5, 95% confidence was performed using Graphpad Prism 5, p < 0.0001.

### *Drosophila* IκBα component, Cactus, is upregulated in the tumor clones

Cact prevents TLR activation by associating with and blocking nuclear localization of Dorsal, the *Drosophila* NF-κB homologue (Dl) or Dorsal-related Immunity Factor (Dif), two effectors of the TLR pathway (16). First, we evaluated Cact expression in tumor cells (16). Cact is ubiquitously expressed, and Cact levels are not affected in wild-type and s*crib^RNAi^* clones (**Fig. 2A, B**, quantified in **Fig. 2E**). However, *Yki^3SA^* and *Yki^3SA^scrib^RNAi^* clones showed significant upregulation of Cact expression (**Fig. 2C, D** quantified in **Fig. 2E**). Likewise, both *Ras^V12^scrib^RNAi^* clones (**Fig. S2A**, quantified in **Fig. S2B**), and *nub>Yki^3SA^scrib^RNAi^* tumors in the wing discs (**Fig. S2C**) showed increased Cact expression. Taken together, our data suggests that tumor cells have increased Cact expression compared to surrounding normal cells. To test if Cact is required for tumorigenesis, we expressed *UAS-cact^RNAi^* and checked if JNK activity is altered when Cact is downregulated in tumor cells. No significant changes in pJNK expression were seen in wild-type (*AyGal4>cact^RNAi^*) (**Fig. 2F**, quantified in **Fig. S2D**) or in *scrib* mutant clones (*AyGal4>cact^RNAi^scrib^RNAi^*) (**Fig. 2G**, quantified in **Fig. S2D**). Interestingly, downregulation of Cact affected cell sorting but not clone size as the jagged clone boundaries in *scrib^RNAi^* clones (**Fig. 2B**), were altered to well-sorted smooth borders in *cact^RNAi^scrib^RNAi^* clones (**Fig. 2G**). In *Yki^3SA^cact^RNAi^* (**Fig. 2H**, quantified in **Fig. S2D**) and *Yki^3SA^scrib^RNAi^cact^RNAi^* clones (**Fig. 2I**, quantified in **J**), downregulation of *cact* prevented pJNK induction, and reduced pJNK expression to wild-type levels. Next, we evaluated the invasiveness of tumor clones when *cact* was downregulated. Downregulation of *cact* in wild-type (**Fig. 2K**) or *scrib^RNAi^* (**Fig. 2L**, quantified in **Fig. S2F**) clones did not affect MMP1 expression. In *Yki^3SA^cact^RNAi^* clones MMP1 levels were significantly reduced (**Fig. 2M**) when compared to *Yki^3SA^* clones, but were significantly higher than wild-type clones (**Fig. S2F, G**). Remarkably, MMP1 levels are reduced to wild-type in *Yki^3SA^ scrib^RNAi^cact^RNAi^* clones (**Fig. 2N**, quantified in **O**) although these clones may still be multi-layered (**Fig. S2E**). Overall, our data showed that decreasing levels of *cact* in the tumor cells reduced JNK activation and MMP1 induction, thereby affecting tumor growth and invasiveness. Interestingly, *nub>Yki^3SA^scrib^RNAi^cact^RNAi^* wing discs also showed reduced pJNK (**Fig. S2H**) and MMP1 (**Fig. S2I**) expression when compared to *nub>Yki^3SA^scrib^RNAi^* wing discs. The difference in pJNK and MMP1 reduction can be attributed to inherent differences in tumor induction mechanism *viz*., by a short pulse of Flippase expression in the ‘flp-out’ clones versus sustained Gal4 driven expression. Overall, these data suggest that elevated levels of *cact* play a key role in tumorigenesis, and downregulation of *cact* affects tumor growth by downregulating JNK signaling.

**Figure 2:**
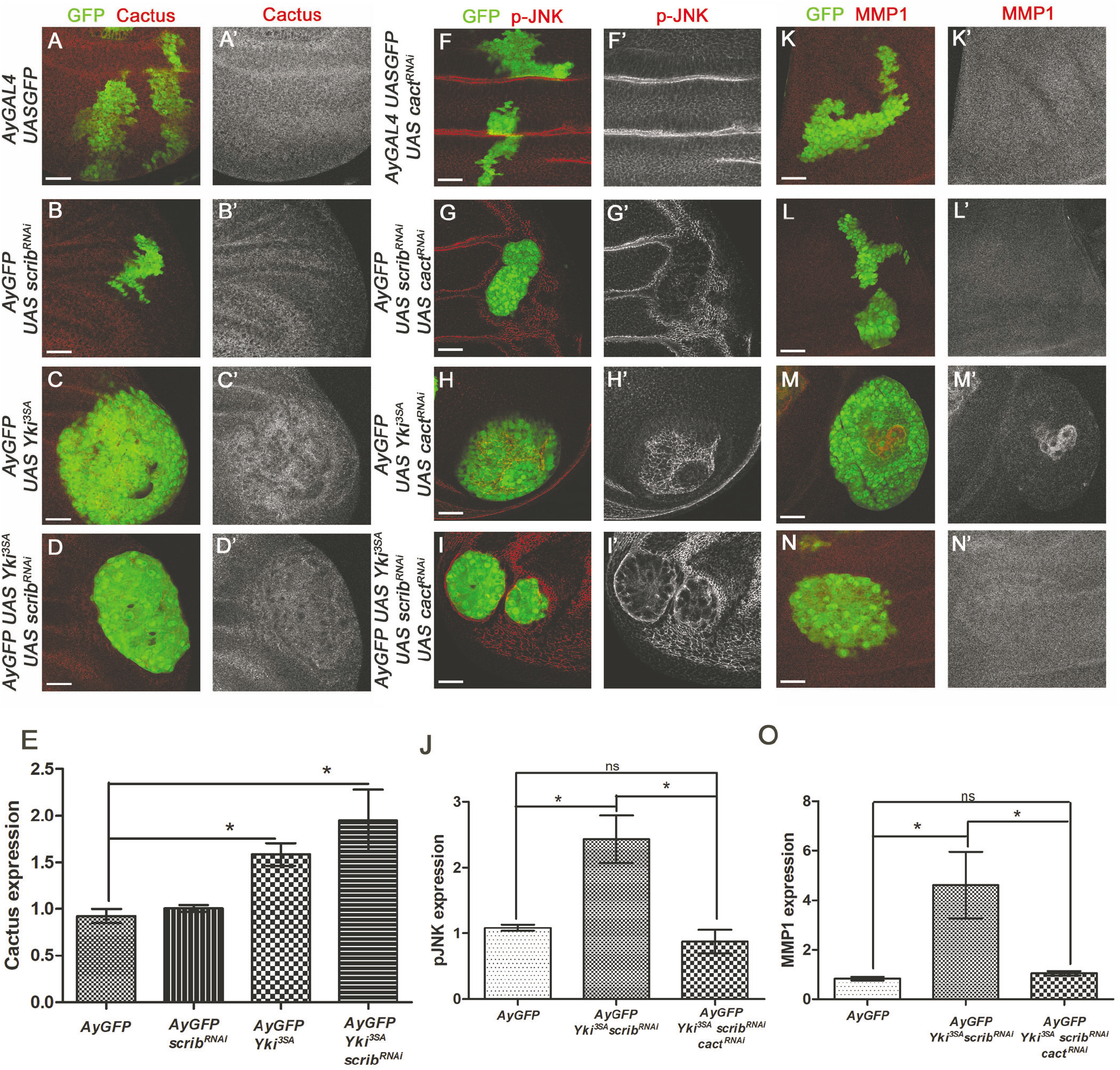
Role of Cact in tumor progression. Panels show Cact expression (red, grey) in GFP-labelled clones (Green) from (**A**) *AyGal4>GFP*, (**B**) *AyGal4>scrib^RNAi^*, (**C**) *AyGAL4>Yki^3SA^* and, (**D**) *AyGAL4> Yki^3SA^scrib^RNAi^* wing discs. (**F-I**) pJNK (red, grey) and (**K-N**) MMP1 expression (red, grey) in clones (green) when Cact is downregulated. (**E, J, O**) Bar graphs depict quantification of change in Cact (**E**), pJNK (**J**) and MMP1 (**O**) expression between indicated genotypes. *= p<0.05, ns= not significant

### TNF pathway affects tumor growth but not invasiveness

Upregulation of the *Drosophila* TNF ligand Egr is shown to induce JNK activity in *Ras^V12^scrib^RNAi^* tumors (5, 6). Egr requires TNF receptors Wgn and /or Grnd for signaling, and downregulation of *wgn* can attenuate Egr activity (31, 32). Hence to manipulate Egr activity in tumor clones, we first downregulated *wgn* through RNA interference in the *Yki^3SA^scrib^RNAi^*. No significant change in pJNK(**Fig. 3A**, quantified in **E)** or MMP1 (**Fig. 3F**, quantified in **J)** expression was observed in clones expressing *wgn^RNAi^*, or *scrib^RNAi^wgn^RNAi^* (**Fig. 3B, G**). However, the size of *scrib^RNAi^wgn^RNAi^* clones was significantly smaller than *scrib^RNAi^* clones (**Fig. S3A**). This is consistent with previous reports where *scrib,egr* double mutant clones grew poorly compared to *scrib* mutant clones (7). In *Yki^3SA^wgn^RNAi^* clones pJNK upregulation is blocked (**Fig. 3C**, quantified in **Fig. S3B**). However, compared to *Yki^3SA^* clones, MMP1 was robustly induced in *Yki^3SA^wgn^RNAi^* clones (**Fig. 3H**, quantified in **Fig. S3C, D**). Similarly, in *Yki^3SA^scrib^RNAi^wgn^RNAi^* clones pJNK activation is suppressed (**Fig. 3D**, quantified in **E**). Interestingly, MMP1 is significantly upregulated in *Yki^3SA^scrib^RNAi^wgn^RNAi^* clones comparable to levels in *Yki^3SA^scrib^RNAi^* clones (**Fig. 3I**, quantified in **Fig. 3J**). Similar findings were reported in *Ras^V12^scrib^RNAi^wgn^RNAi^* tumors or chromosomal instability induced tumor models where MMP1 induction is not affected by downregulation of *wgn* (17, 33). These data suggested that either the wild-type levels of pJNK are sufficient for MMP1 induction in tumors, or Grnd, the other TNF receptor, regulates the activation of TNF-JNK pathway during tumor growth and invasion.

**Figure 3:**
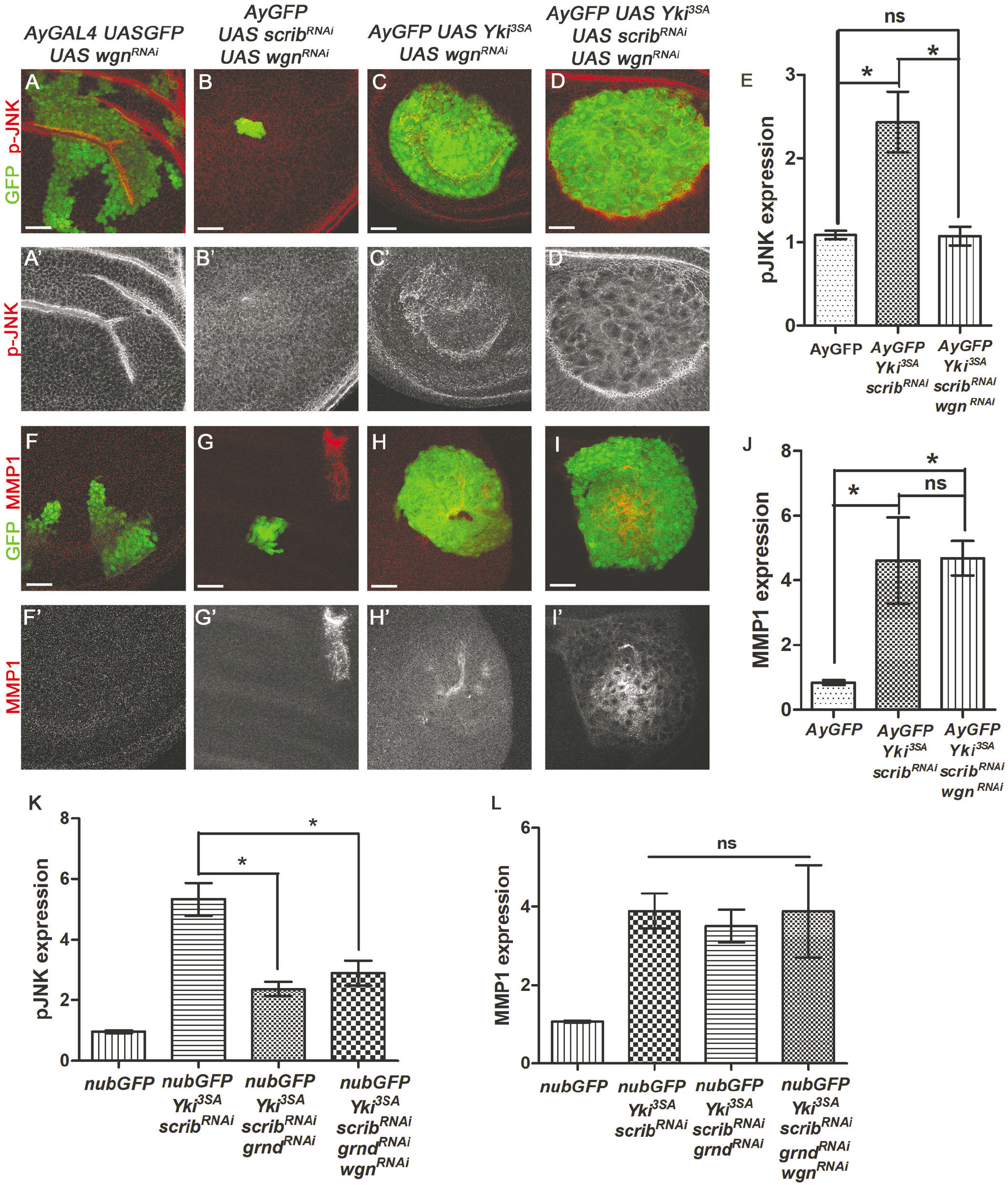
Effect of downregulating both the TNF receptors on the Tumor progression. Panels show wing imaginal discs containing heat-shock induced GFP labelled (green) clones expressing *wgn^RNAi^* in the indicated genotypes stained with Rb anti-pJNK antibody (**A-D**) or anti-MMP1 antibody (**G-J**). (**E, J-L**) Bar graphs showing quantification of pJNK (**E, K**) and MMP1 (**F, L**) expression amongst indicated genotypes. *= p<0.05, ns= not significant

We could not generate *Act>y+>Gal4UASgrnd^RNAi^* recombined lines since ‘flp-out’ clones made with *eyFLP* or *hsFLP* lacked a definitive phenotype. However, we succeeded in recombining *nubGal4 UASGFP* with *UASgrnd^RNAi^* flies (details in supplementary methods). Similar to *wgn^RNAi^*, pJNK levels were significantly reduced in wing discs from *Yki^3SA^scrib^RNAi^grnd^RNAi^* larvae when compared to *Yki^3SA^scrib^RNAi^*, but significantly higher than wild-type (*nub>GFP*) levels (**Fig. S4A**, quantified in **Fig. 3K**). However, MMP1 was still significantly induced (**Fig. S4B** quantified in **Fig. 3L**). Previous reports show that tumors caused by chromosomal instability show high MMP1 induction despite *grnd* downregulation (33). To account for the possible functional redundancy between Wgn and Grnd, we downregulated both receptors and evaluated MMP1 expression in *Yki^3SA^scrib^RNAi^* tumors. Compared to *nub>Yki^3SA^scrib^RNAi^*, downregulation of both TNF receptors (*nub>Yki^3SA^scribi^RNAi^grnd^RNAi^wgn^RNAi^*) reduced pJNK levels significantly (**Fig. S4C**, quantified in **Fig. 3K**), but no significant reduction in MMP1 expression was observed (**Fig. S4D**, quantified in **Fig. 3L**). Taken together, these data suggest that tumor growth but not invasiveness is regulated by TNF pathway receptors.

### JNK regulates invasiveness but not Cact expression in the tumor

Although downregulating TNF receptors didn’t affect MMP1 induction in the tumor cells (**Fig. 3K, L**), downregulating TLR component, Cact, reduced MMP1 expression to wild-type levels (**Fig. 2O**). Given that JNK is the key MAPK that can transcriptionally activate MMP1(27), we investigated if Cact can intracellularly activate JNK independent of the TNF receptors and cause MMP1 activation in tumor cells. We compared pJNK and MMP1 expression in the *Yki^3SA^scrib^RNAi^ wgn^RNAi^* clones when JNK activity was inhibited by expressing dominant negative form of *Drosophila* JNK, *bsk^DN^* (34). As expected, downregulation of JNK in the *Yki^3SA^ scrib^RNAi^* clones (*Yki^3SA^scrib^RNAi^bsk^DN^*) inhibited both pJNK activation (**Fig. 4A**, quantified in **D**) and MMP1 induction (**Fig. 4E**, quantified in **G**). Interestingly, co-expression of *bsk^DN^* with *Yki^3SA^scrib^RNAi^wgn^RNAi^*, blocked both pJNK (**Fig. 4B**, quantified in **D**) and MMP1 induction (**Fig. 4F** quantified in **G**). Thus, in tumor cells MMP1 expression can be stimulated by a TNF receptor-independent intracellular JNK activation mechanism. Furthermore, downregulation of JNK and Wgn significantly reduced clone thickness (15.63μm) (**Fig. 4C**) as compared to that of *Yki^3SA^scrib^RNAi^* tumors (23.17μm). Interestingly while Cact downregulation affected JNK activation, downregulating JNK in tumor clones (*bsk^DN^Yki^3SA^scrib^RNAi^*) showed significant Cact expression similar to *Yki^3SA^scrib^RNAi^* tumor clones (**Fig. 4H**, quantified in **I**). Thus our data showed, Cact acts upstream of JNK signaling, and affects oncogenic JNK signaling.

**Figure 4:**
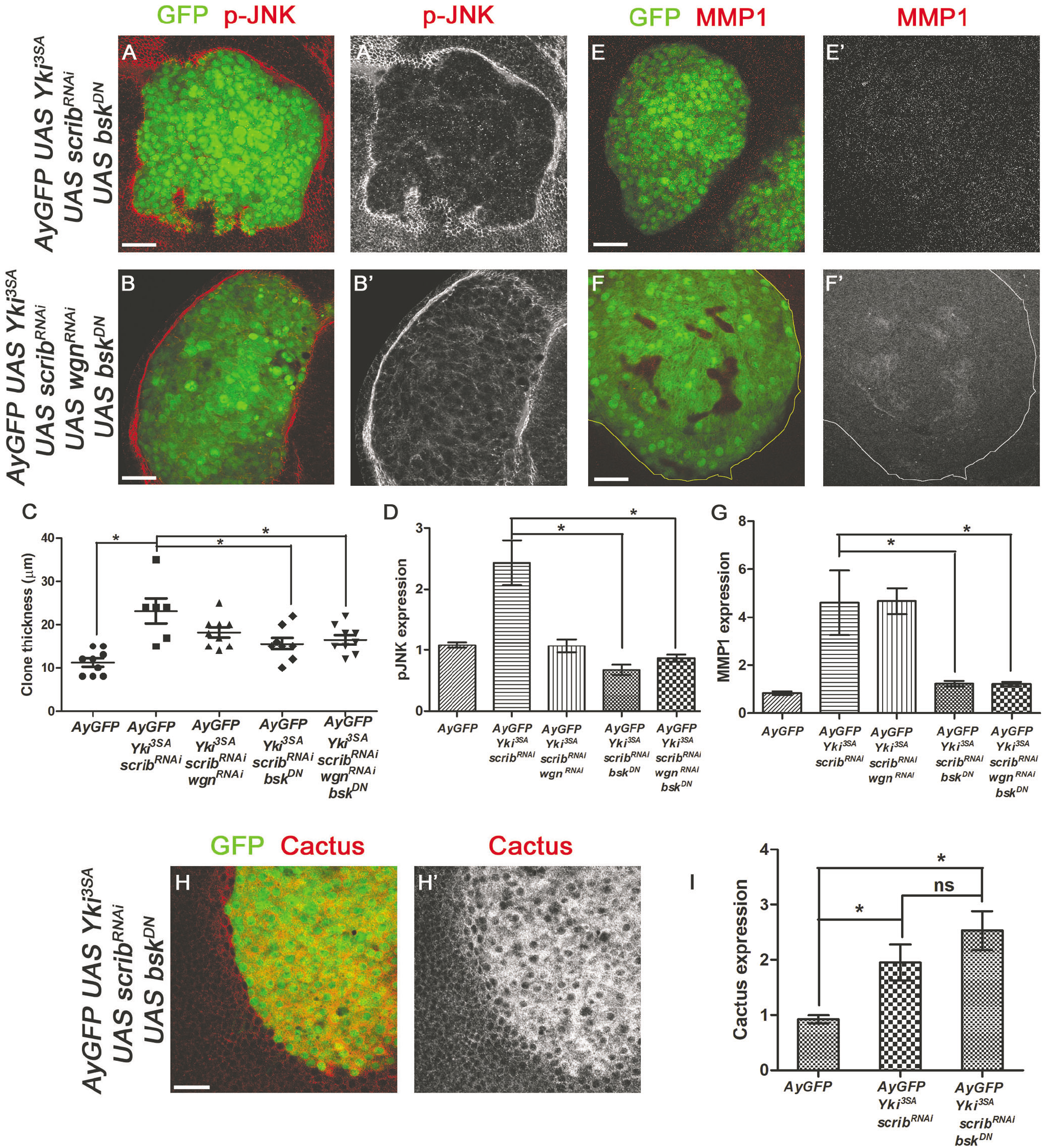
JNK is the key regulator of tumor invasiveness. Panels (**A-B’, E-F’, H-H’**) show GFP-labelled heat-shock clones (green) in the wing imaginal discs of *Yki^3SA^scrib^RNAi^bsk^DN^* (**A, E, H**) and *Yki^3SA^scrib^RNAi^bsk^DN^wgn^RNAi^* (**B, F**) stained for antibodies against pJNK (red, grey) (**A, B**), MMP1 (red, grey) (**E, F**), and Cact (red, grey) (**H**). (**C**) Thickness of the clones representing the multi-layered structure of indicated genotypes was measured using Photoshop, n=8. (**D, G, I**) Bar graphs show quantification of change in expression of pJNK (**D**), MMP1 (**G**) and Cact (**I**) amongst indicated genotype. *= p<0.05, ns= not significant

### Yki regulates Cact and JNK in promoting tumor survival and invasiveness

Oncogenic cooperation can induce Yki and JNK activity in a context-dependent manner (35, 36), and during anti-microbial response Yki can transcriptionally regulate Cact expression (37). Given this, we investigated the role of Yki in regulating Cact and JNK levels in our tumor models. We downregulated the levels of Yki (*UASyki^RNAi^*) in *Ras^V12^scrib^RNAi^* tumor clones. Under our experimental condition (5min heat shock at 37°C to second instar larvae), clones expressing *yki^RNAi^* were competed out when the larvae were grown at 25 °C or 18 °C. Therefore, to prevent elimination of *yki^RNAi^* clones, we co-expressed p35- a pan caspase inhibitor for our experiments (38). Co-expression of *p35* and *yki^RNAi^* didn’t affect Cact expression in the wild-type (**Fig. 5A**), *scrib^RNAi^* (**Fig. 5B**), or *Ras^V12^* clones (**Fig. 5C**). In comparison to control clones (**Fig. 5A-C**), the *Ras^V12^scrib^RNAi^yki^RNAi^* clones survived, but showed significant reduction in both the clone size (**Fig. 5E**) and thickness (**Fig. 5F**). Interestingly, in *Ras^V12^scrib^RNAi^yki^RNAi^* clones downregulation of Yki blocked Cact (**Fig. 5D**, quantified in **G**), pJNK (**Fig. S5A**) and MMP1(**Fig. S5B**) induction as compared to *Ras^V12^scrib^RNAi^* clones.

**Figure 5:**
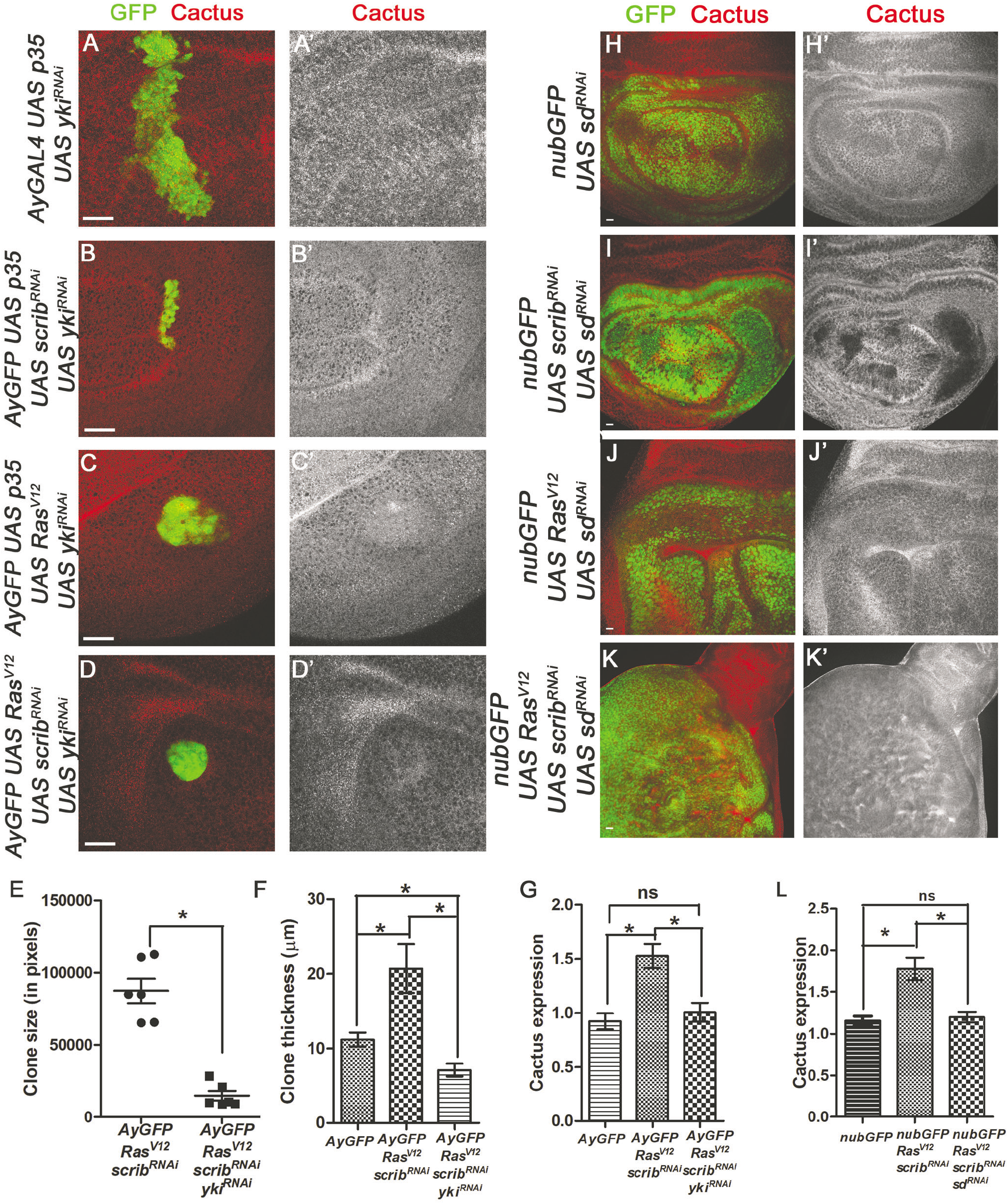
Role of Yki in stimulating Cact and promoting tumorigenesis. Panels **A-F** show effect of downregulating *yki* (*UASyki^RNAi^*) in the GFP-labelled heat-shock clones (green) in wing imaginal discs from larvae of the indicated genotype. (**A-D**) Confocal images present effect of *yki^RNAi^* on Cact (red, grey) expression. (**E-G**) Graphs show quantification of clone size (**E**), clone thickness (**F**), and change in Cact expression levels (**G**). Panels **H-L** show effects of *sd* downregulation (*UASsd^RNAi^*) on *nubGal4* driven tumor phenotypes in the wing imaginal discs. (**H-K**) Confocal images of wing discs from indicated genotypes stained with anti-Cact antibody (red, grey) are presented. (**L**) Graph showing quantification of changes in Cact expression. *= p<0.05, ns= not significant

The TEAD family transcription factor Scalloped (Sd) is the major binding partner for Yki mediated transcription (39–41). The Yki/Sd complex also regulates *cact* expression during innate immune response (37). Sd downregulation (*nub>sd^RNAi^*) resulted in a diminished wing pouch and yielded adult flies with smaller wings (42). Cact levels are not affected in wing discs from *nub>sd^RNAi^* (**Fig. 5H**) or *nub>scrib^RNAi^sd^RNAi^* larvae (**Fig. 5I**). High larval lethality was seen in *nub>Ras^V12^sd^RNAi^* and *nub>Ras^V12^scrib^RNAi^sd^RNAi^* larvae grown at 25°C. However, at 18°C, Cact was not induced in *Ras^V12^sd^RNAi^* (**Fig. 5J**) and *Ras^V12^scrib^RNAi^sd^RNAi^* wing discs (**Fig. 5K**, quantified in **L**). In addition, pJNK and MMP1 expression were also not induced in *Ras^V12^scrib^RNAi^sd^RNAi^* wing pouch (Fig. **S5C, D**). These results strongly support a model where Yki and Sd regulate Cact expression, and loss of either Yki or Sd is sufficient to prevent the upregulation of Cact in the tumor cells. Further downregulation of Yki, Sd or Cact is sufficient to cause downregulation of JNK-mediated sustained signaling that promotes proliferation and invasion. These results suggest a mechanism in which polarity deficient cells with activated Ras require Yki mediated Cact induction and JNK activation to drive tumor growth and invasiveness (**Fig. 6**).

**Figure 6:**
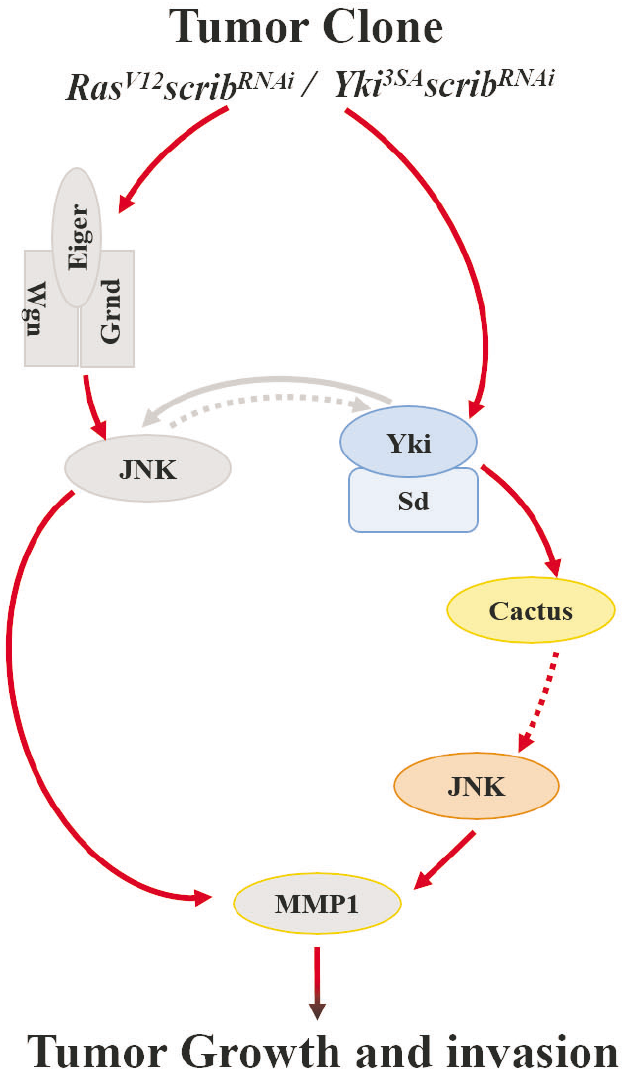
Model for Yki mediated Cact and JNK activation promoting tumorigenesis. A model depicting the downstream effects of the cooperation between activated oncogene (*Yki* or *Ras*) and loss of polarity gene, *scrib* is shown. In somatic clones (elliptical cells), JNK is activated by TNF pathway ligand Egr and its receptors, Wgn and Grnd. Independently, JNK can be activated in the tumor cells due to Cact accumulation via Yki activation. Yki mediated Cact and JNK activation is critical for tumor progression.

## Discussion

### Yki dependent signals downstream of Oncogenic Ras promote tumor growth

Oncogenic Ras activation results in burst of signaling through downstream effectors (Raf, Rac, Rho, PI3K), which phosphorylate and activate transcription factors, such as c-Myc, c-Jun, and c-Fos (27, 43). The plethora of downstream signals make it difficult to identify and analyze key signaling nodes that promote tumor growth. Recently, Hippo pathway effectors YAP/Yki have emerged as key effectors of activated Ras signaling (11, 12, 14, 15). Moreover, YAP/Yki are sites of signal convergence and integration in many mammalian cancers e.g., colon, pancreatic and lung cancer (13, 44). The JNK pathway has emerged as another tumor promoting mechanism that can exert both anti- and pro-tumor activities (24, 45). JNK signaling induces Yki activation during compensatory cell proliferation and neoplastic tumor growth (7, 36, 46, 47), and JNK suppresses Yki elevation in *scrib* mutant cells during growth regulation (48, 49). Thus, JNK, Yki, and their downstream transcription factors have emerged as synergistic drivers of tumor growth (35, 50–53).

Consistent with previous reports, overexpression of *Yki^3SA^* alone leads to hyperplasia, and homozygous loss of *scrib* induces neoplasia (42, 54, 55). When *Yki^3SA^scrib^RNAi^* are co-expressed, multilayered metastatic tumors formed that degrade the basement membrane (**Fig. 1**). We confirmed that JNK signaling is activated in *Yki^3SA^scrib^-^* induced neoplastic growth (**Fig. 1**). Polyploid giant cells in *Ras^V12^scrib^-/-^* clones are linked to increased Yki and JNK activity that promotes tumor progression (56). Variation in cell size is usually seen in primary tumor cells whereas larger metastases show more uniform size (57). We found cells of variable size in the *Yki^3SA^scrib^RNAi^* clones (**Fig.1C**) but uniform sized in *nub>Yki^3SA^scrib^RNAi^* discs. Further, impaired Hippo signaling can induce JNK signaling via transcriptional activation of the Rho1-GTPase which mediates Yki-induced JNK activation and overgrowth (58). Similarly, in mammals, GPCRs can activate YAP/TAZ through RhoA suggesting Yki-Rho1-JNK axis involvement in bridging the two pathways (58, 59). Our data and other published work suggests that both Yki and JNK dependent signaling is activated in multiple tumor contexts including *Yki^3SA^scrib^RNAi^* tumors. A key transcriptional target of JNK signaling is MMP1- a matrix metalloprotease required for tissue remodeling, wound healing, regeneration, and control of cell movement and cell adhesion during development; and degradation of the basement membrane during tumor invasion (25, 27–29, 60, 61). We used JNK activation and MMP1 induction as the criteria to evaluate tumor growth and invasiveness. The *Yki^3SA^scrib^RNAi^* tumor clones showed high levels of JNK activity and MMP1 induction (**Fig. 1**) suggesting that the tumor cells have acquired invasive traits.

### Yki regulates Cact and JNK in Tumor cells

Cact is a negative regulator of the *Drosophila* Rel/NFκB orthologues Dorsal, Dif and Relish. Degradation of Cact, allows NFκB activation and localization to nucleus where it regulates transcription of genes controlling inflammatory cytokines and cell growth (62, 63). However, we noticed an accumulation of Cact in our tumor clones. (**Fig. 2**). Downregulating Cact in tumor cells prevented tumor progression by suppressing JNK activation and MMP1 induction (**Fig. 2**). This indicated a role of Cact in promoting tumor progression. Increase in Toll signaling by downregulation of Cact aggravates JNK activated cell death (64). Interestingly, we observed a significant reduction in pJNK expression with downregulation of Cact in our tumor clones (**Fig. 2**). These observations indicate that IκBα can regulate proliferation through JNK. Moreover, we found that downregulating JNK by downregulating TNF receptors (**Fig. 3**) did not affect MMP1 induction in our tumor clone. Also, downregulation of JNK by expressing dominant negative form of JNK (**Fig. 4**), did not affect Cact expression in the tumor cells although it blocked JNK and MMP1 activation (**Fig. 4**). Based on these genetic interactions, we concluded that Cact acts upstream of JNK and plays a key role in regulation of oncogenic JNK signaling (**Fig. 6**). We also observed that Wgn downregulation in tumor clones did not affect Cact accumulation (data not shown). Although JNK levels were reduced, MMP1 was still induced in such cells (**Fig. 3**). Since TNFR1 can antagonistically regulate cell cycle through JNK and NFκB (65), it will be interesting to explore whether such signaling crosstalk between these three pathways affects mammalian tumorigenesis.

JNK is known to play a context dependent role to promote tumor growth. Work from our lab and others found that, Yki can also regulate JNK activation in the tumor cells. Liu et al. demonstrated that Yki can regulate the TLR pathway and prevent antimicrobial response (37). They showed that in the *Drosophila* larval fat body, Yki along with its transcriptional co-activator Scalloped, is capable of activating Cact. We also observed that downregulating either Yki or Scalloped severely affected tumor growth and showed no overexpression of Cact (**Fig. 5**). Further, downregulating Yki in tumor cells also restored JNK level. Similarly, in Basal Cell Carcinoma YAP interacts with its DNA binding transcription factors, TEAD and promotes c-Jun activity while loss of YAP leads to reduction in phosphorylated JNK and JUN (66). Overall, based on these data we propose a new mechanism where Yki functions with Sd to activate Cact and regulate JNK induction in the tumor cells (**Fig. 6**). Although *cact* is a transcriptional target of the Yki/Sd complex during innate immune response (37), whether this interaction is involved in tumor immunogenicity remains unknown. Recent studies have shown that inactivation of Hippo pathway in tumor cells induces host inflammatory responses, and the pathway can respond to and mediate inflammatory signals (67, 68). In conclusion, our study unraveled the interaction between the Yki, JNK and Cact to drive tumorigenesis. Therefore, it will be interesting to explore the mechanisms underlying the role of innate immunity in tumor progression and metastasis.

## Material and methods

### Fly Strains and Generation of Clones

All fly mutant and transgenic lines are described in Flybase, and were obtained from the Bloomington Drosophila Stock Center unless otherwise specified. *UAS-GFP* labeled clones were produced in larval imaginal discs using the following strains: *nub-Gal4/CyO* (from S. Cohen)-, *yw; Act>y+>Gal4,UAS-GFP* (BL4411), *UASYki^3SA^*(BL28817), *UASscrib^RNAi^* (BL58085, BL59080), *UASwgn^RNAi^* (BL50594), *UASgrnd^RNAi^/CyO* (from A. Bergmann), *UAScact^RNAi^* (BL31713), *UASRas^V12^* (BL5788), *UASbsk^DN^* (BL6409), *UASyki^RNAi^* (BL34067), *UASsd^RNAi^*(BL29352),*UASP35*, and *puc*^E69^-*lacZ*. Appropriate genetic crosses were performed to establish recombined fly stocks to generate clones of the indicated genotypes. To induce somatic clones, larvae (48 hr after egg laying) were heat shocked at 37°C for 5 min. This heat shock regimen was followed to induce *Yki^3SA^ scrib^RNAi^* clones alone or in combination with other transgenes described in the manuscript.

### Immunohistochemistry

Third instar larvae were dissected and processed for immunohistochemistry following standard protocol (69). The samples were mounted in the VectaShield mounting medium (Vector Labs) and scanned using confocal microscopy (Olympus Fluoview 1000, 3000). The primary antibodies used were mouse anti-Cact (DHSB, 1:200), rabbit anti-pJNK (Cell Signaling Technology, 1:250), mouse anti-β gal (DSHB, 1:250), rabbit anti-Laminin (Ab-Cam, 1:250) and mouse anti-MMP1 (DHSB, 1:200). Secondary antibody used were donkey Cy3-conjugated anti-mouse IgG (1:200, Jackson ImmunoResearch), and donkey Cy3-conjugated anti-rabbit IgG (1:200, Jackson ImmunoResearch).

### Quantification

The immunohistochemistry data was quantified using the measurement log function of Photoshop (Adobe Photoshop CC 2018). For the heat shock induced clones, 3 circular ROI of 50pixel radius were used per clone (GFP positive) and adjacent normal region (GFP-negative). For the *nubGAL4* driven experiments, 3 square ROIs of 100 pixel length were used. The average value of the integrated density was compared in between the normal non-GFP and clone specific GFP positive cells. For comparison between two different genotypes, ratio of the average integrated density in the GFP positive and negative region was compared. A ratio of 1 indicated no change in expression level between the tumor (GFP positive) and normal (GFP negative) region. In all studies, the change in expression compared between different genotypes is normalized with wild-type levels set to 1. Thickness of the clones was measured using the inbuilt ruler of the Fluoview 3000 software in the XZ/YZ optical sections for all experiments. Statistical significance was quantified by Student t-test using Graphpad Prism 5 software.

## Acknowledgements

We would like to thank, Dr. A. Bergmann, Dr. Stephen Cohen, the Bloomington Drosophila Stock Center, and the Drosophila Studies Hybridoma Bank for flies and antibodies. KS acknowledge the Teaching Assistantship and Graduate Student Summer Fellowship (GSSF) from the Graduate Program of University of Dayton. AS is supported by Start-up research funds and funding from Schuellein Endowed Chair in Biology from the University of Dayton and NIH 1R15 GM124654-1. MKS is supported by start-up research funds from the University of Dayton, and a subaward from NIH grant R01CA183991 (PI Nakano).

## Conflict of Interest

The authors declare no conflict of interest.

**Supplementary information is available at Oncogene’s website**.

## References

1. Fernandez-Medarde A, Santos E. Ras in cancer and developmental diseases. Genes Cancer. 2011;2(3):344–58.

2. Young A, Lou D, McCormick F. Oncogenic and wild-type Ras play divergent roles in the regulation of mitogen-activated protein kinase signaling. Cancer Discov. 2013;3(1):112–23.

3. Pagliarini RA, Xu T. A genetic screen in Drosophila for metastatic behavior. Science. 2003;302(5648):1227–31.

4. Brumby AM, Richardson HE. scribble mutants cooperate with oncogenic Ras or Notch to cause neoplastic overgrowth in Drosophila. EMBO J. 2003;22(21):5769–79.

5. Cordero JB, Macagno JP, Stefanatos RK, Strathdee KE, Cagan RL, Vidal M. Oncogenic Ras diverts a host TNF tumor suppressor activity into tumor promoter. Dev Cell. 2010;18(6):999–1011.

6. Igaki T, Kanda H, Yamamoto-Goto Y, Kanuka H, Kuranaga E, Aigaki T, et al. Eiger, a TNF superfamily ligand that triggers the Drosophila JNK pathway. EMBO J. 2002;21(12):3009–18.

7. Ohsawa S, Sugimura K, Takino K, Xu T, Miyawaki A, Igaki T. Elimination of oncogenic neighbors by JNK-mediated engulfment in Drosophila. Dev Cell. 2011;20(3):315–28.

8. Hanahan D, Weinberg RA. Hallmarks of cancer: the next generation. Cell. 2011;144(5):646–74.

9. Mantovani A, Allavena P, Sica A, Balkwill F. Cancer-related inflammation. Nature. 2008;454(7203):436–44.

10. Shao DD, Xue W, Krall EB, Bhutkar A, Piccioni F, Wang X, et al. KRAS and YAP1 converge to regulate EMT and tumor survival. Cell. 2014;158(1):171–84.

11. Kapoor A, Yao W, Ying H, Hua S, Liewen A, Wang Q, et al. Yap1 activation enables bypass of oncogenic Kras addiction in pancreatic cancer. Cell. 2014;158(1):185–97.

12. Snigdha K, Gangwani KS, Lapalikar GV, Singh A, Kango-Singh M. Hippo Signaling in Cancer: Lessons From Drosophila Models. Frontiers in Cell and Developmental Biology. 2019;7(85).

13. Zanconato F, Cordenonsi M, Piccolo S. YAP/TAZ at the Roots of Cancer. Cancer Cell. 2016;29(6):783–803.

14. Zhang W, Nandakumar N, Shi Y, Manzano M, Smith A, Graham G, et al. Downstream of mutant KRAS, the transcription regulator YAP is essential for neoplastic progression to pancreatic ductal adenocarcinoma. Sci Signal. 2014;7(324):ra42.

15. Mao Y, Sun S, Irvine KD. Role and regulation of Yap in KrasG12D-induced lung cancer. Oncotarget. 2017;8(67):110877–89.

16. Valanne S, Wang JH, Ramet M. The Drosophila Toll signaling pathway. J Immunol. 2011;186(2):649–56.

17. Andersen DS, Colombani J, Palmerini V, Chakrabandhu K, Boone E, Rothlisberger M, et al. The Drosophila TNF receptor Grindelwald couples loss of cell polarity and neoplastic growth. Nature. 2015;522(7557):482–6.

18. Calleja M, Moreno E, Pelaz S, Morata G. Visualization of gene expression in living adult Drosophila. Science. 1996;274(5285):252–5.

19. Birembaut P, Caron Y, Adnet JJ, Foidart JM. Usefulness of basement membrane markers in tumoural pathology. J Pathol. 1985;145(4):283–96.

20. Ito K, Awano W, Suzuki K, Hiromi Y, Yamamoto D. The Drosophila mushroom body is a quadruple structure of clonal units each of which contains a virtually identical set of neurones and glial cells. Development. 1997;124(4):761–71.

21. Kango-Singh M, Singh A, Henry Sun Y. Eyeless collaborates with Hedgehog and Decapentaplegic signaling in Drosophila eye induction. Dev Biol. 2003;256(1):49–60.

22. Dahmann C, Basler K. Opposing transcriptional outputs of Hedgehog signaling and engrailed control compartmental cell sorting at the Drosophila A/P boundary. Cell. 2000;100(4):411–22.

23. Cellurale C, Sabio G, Kennedy NJ, Das M, Barlow M, Sandy P, et al. Requirement of c-Jun NH(2)-terminal kinase for Ras-initiated tumor formation. Mol Cell Biol. 2011;31(7):1565–76.

24. Uhlirova M, Jasper H, Bohmann D. Non-cell-autonomous induction of tissue overgrowth by JNK/Ras cooperation in a Drosophila tumor model. Proc Natl Acad Sci U S A. 2005;102(37):13123–8.

25. Igaki T, Pagliarini RA, Xu T. Loss of cell polarity drives tumor growth and invasion through JNK activation in Drosophila. Curr Biol. 2006;16(11):1139–46.

26. Martin-Blanco E, Gampel A, Ring J, Virdee K, Kirov N, Tolkovsky AM, et al. puckered encodes a phosphatase that mediates a feedback loop regulating JNK activity during dorsal closure in Drosophila. Genes Dev. 1998;12(4):557–70.

27. Uhlirova M, Bohmann D. JNK- and Fos-regulated Mmp1 expression cooperates with Ras to induce invasive tumors in Drosophila. EMBO J. 2006;25(22):5294–304.

28. Deryugina EI, Quigley JP. Matrix metalloproteinases and tumor metastasis. Cancer Metastasis Rev. 2006;25(1):9–34.

29. Beaucher M, Hersperger E, Page-McCaw A, Shearn A. Metastatic ability of Drosophila tumors depends on MMP activity. Dev Biol. 2007;303(2):625–34.

30. Page-McCaw A, Serano J, Sante JM, Rubin GM. Drosophila matrix metalloproteinases are required for tissue remodeling, but not embryonic development. Dev Cell. 2003;4(1):95–106.

31. Kauppila S, Maaty WS, Chen P, Tomar RS, Eby MT, Chapo J, et al. Eiger and its receptor, Wengen, comprise a TNF-like system in Drosophila. Oncogene. 2003;22(31):4860–7.

32. Kanda H, Igaki T, Kanuka H, Yagi T, Miura M. Wengen, a member of the Drosophila tumor necrosis factor receptor superfamily, is required for Eiger signaling. J Biol Chem. 2002;277(32):28372–5.

33. Muzzopappa M, Murcia L, Milan M. Feedback amplification loop drives malignant growth in epithelial tissues. Proc Natl Acad Sci U S A. 2017;114(35):E7291–E300.

34. Willsey HR, Zheng X, Carlos Pastor-Pareja J, Willsey AJ, Beachy PA, Xu T. Localized JNK signaling regulates organ size during development. Elife. 2016;5.

35. Atkins M, Potier D, Romanelli L, Jacobs J, Mach J, Hamaratoglu F, et al. An Ectopic Network of Transcription Factors Regulated by Hippo Signaling Drives Growth and Invasion of a Malignant Tumor Model. Curr Biol. 2016;26(16):2101–13.

36. Enomoto M, Kizawa D, Ohsawa S, Igaki T. JNK signaling is converted from anti- to pro-tumor pathway by Ras-mediated switch of Warts activity. Dev Biol. 2015;403(2):162–71.

37. Liu B, Zheng Y, Yin F, Yu J, Silverman N, Pan D. Toll Receptor-Mediated Hippo Signaling Controls Innate Immunity in Drosophila. Cell. 2016;164(3):406–19.

38. Hay BA, Wolff T, Rubin GM. Expression of baculovirus P35 prevents cell death in Drosophila. Development. 1994;120(8):2121–9.

39. Goulev Y, Fauny JD, Gonzalez-Marti B, Flagiello D, Silber J, Zider A. SCALLOPED interacts with YORKIE, the nuclear effector of the hippo tumor-suppressor pathway in Drosophila. Curr Biol. 2008;18(6):435–41.

40. Wu S, Liu Y, Zheng Y, Dong J, Pan D. The TEAD/TEF family protein Scalloped mediates transcriptional output of the Hippo growth-regulatory pathway. Dev Cell. 2008;14(3):388–98.

41. Zhang L, Ren F, Zhang Q, Chen Y, Wang B, Jiang J. The TEAD/TEF family of transcription factor Scalloped mediates Hippo signaling in organ size control. Dev Cell. 2008;14(3):377–87.

42. Verghese S, Waghmare I, Kwon H, Hanes K, Kango-Singh M. Scribble acts in the Drosophila fat-hippo pathway to regulate warts activity. PLoS One. 2012;7(11):e47173.

43. Rajalingam K, Schreck R, Rapp UR, Albert S. Ras oncogenes and their downstream targets. Biochim Biophys Acta. 2007;1773(8):1177–95.

44. Liu AM, Xu Z, Luk JM. An update on targeting Hippo-YAP signaling in liver cancer. Expert Opin Ther Targets. 2012;16(3):243–7.

45. Tournier C. The 2 Faces of JNK Signaling in Cancer. Genes Cancer. 2013;4(9-10):397–400.

46. Sun G, Irvine KD. Regulation of Hippo signaling by Jun kinase signaling during compensatory cell proliferation and regeneration, and in neoplastic tumors. Dev Biol. 2011;350(1):139–51.

47. Sun G, Irvine KD. Ajuba family proteins link JNK to Hippo signaling. Sci Signal. 2013;6(292):ra81.

48. Chen CL, Schroeder MC, Kango-Singh M, Tao C, Halder G. Tumor suppression by cell competition through regulation of the Hippo pathway. Proc Natl Acad Sci U S A. 2012;109(2):484–9.

49. Doggett K, Grusche FA, Richardson HE, Brumby AM. Loss of the Drosophila cell polarity regulator Scribbled promotes epithelial tissue overgrowth and cooperation with oncogenic Ras-Raf through impaired Hippo pathway signaling. BMC Dev Biol. 2011;11:57.

50. Bunker BD, Nellimoottil TT, Boileau RM, Classen AK, Bilder D. The transcriptional response to tumorigenic polarity loss in Drosophila. Elife. 2015;4.

51. Zanconato F, Forcato M, Battilana G, Azzolin L, Quaranta E, Bodega B, et al. Genome-wide association between YAP/TAZ/TEAD and AP-1 at enhancers drives oncogenic growth. Nat Cell Biol. 2015;17(9):1218–27.

52. Patel PH, Dutta D, Edgar BA. Niche appropriation by Drosophila intestinal stem cell tumours. Nat Cell Biol. 2015;17(9):1182–92.

53. Kulshammer E, Mundorf J, Kilinc M, Frommolt P, Wagle P, Uhlirova M. Interplay among Drosophila transcription factors Ets21c, Fos and Ftz-F1 drives JNK-mediated tumor malignancy. Dis Model Mech. 2015;8(10):1279–93.

54. Oh H, Reddy BV, Irvine KD. Phosphorylation-independent repression of Yorkie in Fat-Hippo signaling. Dev Biol. 2009;335(1):188–97.

55. Morimoto K, Tamori Y. Induction and Diagnosis of Tumors in Drosophila Imaginal Disc Epithelia. J Vis Exp. 2017(125).

56. Cong B, Ohsawa S, Igaki T. JNK and Yorkie drive tumor progression by generating polyploid giant cells in Drosophila. Oncogene. 2018;37(23):3088–97.

57. Bell CD, Waizbard E. Variability of cell size in primary and metastatic human breast carcinoma. Invasion Metastasis. 1986;6(1):11–20.

58. Ma X, Chen Y, Xu W, Wu N, Li M, Cao Y, et al. Impaired Hippo signaling promotes Rho1-JNK-dependent growth. Proc Natl Acad Sci U S A. 2015;112(4):1065–70.

59. Yu FX, Zhao B, Panupinthu N, Jewell JL, Lian I, Wang LH, et al. Regulation of the Hippo-YAP pathway by G-protein-coupled receptor signaling. Cell. 2012;150(4):780–91.

60. Ramet M, Lanot R, Zachary D, Manfruelli P. JNK signaling pathway is required for efficient wound healing in Drosophila. Dev Biol. 2002;241(1):145–56.

61. Page-McCaw A, Ewald AJ, Werb Z. Matrix metalloproteinases and the regulation of tissue remodelling. Nat Rev Mol Cell Biol. 2007;8(3):221–33.

62. Baldwin AS, Jr. The NF-kappa B and I kappa B proteins: new discoveries and insights. Annu Rev Immunol. 1996;14:649–83.

63. Xia Y, Shen S, Verma IM. NF-kappaB, an active player in human cancers. Cancer Immunol Res. 2014;2(9):823–30.

64. Wu C, Chen C, Dai J, Zhang F, Chen Y, Li W, et al. Toll pathway modulates TNF-induced JNK-dependent cell death in Drosophila. Open Biol. 2015;5(7):140171.

65. Zhang JY, Tao S, Kimmel R, Khavari PA. CDK4 regulation by TNFR1 and JNK is required for NF-kappaB-mediated epidermal growth control. J Cell Biol. 2005;168(4):561–6.

66. Maglic D, Schlegelmilch K, Dost AF, Panero R, Dill MT, Calogero RA, et al. YAP-TEAD signaling promotes basal cell carcinoma development via a c-JUN/AP1 axis. EMBO J. 2018;37(17).

67. Nowell CS, Odermatt PD, Azzolin L, Hohnel S, Wagner EF, Fantner GE, et al. Chronic inflammation imposes aberrant cell fate in regenerating epithelia through mechanotransduction. Nat Cell Biol. 2016;18(2):168–80.

68. Taniguchi K, Wu LW, Grivennikov SI, de Jong PR, Lian I, Yu FX, et al. A gp130-Src-YAP module links inflammation to epithelial regeneration. Nature. 2015;519(7541):57–62.

69. Kango-Singh M, Nolo R, Tao C, Verstreken P, Hiesinger PR, Bellen HJ, et al. Shar-pei mediates cell proliferation arrest during imaginal disc growth in Drosophila. Development. 2002;129(24):5719–30.

