## Supplementary material and figures for "THE YKI-CACTUS (I_K_Bα)-JNK AXIS PROMOTES TUMOR GROWTH AND PROGRESSION IN DROSOPHILA"

Snigdha et al 2019

### Supplementary Information

**Supplementary Methods:**

**Recombination of *UASgrnd*$^{RNAi}$ with *nubGal4 UASGFP*:** Overexpression of *UASgrnd*$^{RNAi}$ with several GAL4 drivers (including *nubGAL4*) in the wing discs did not produce a phenotype in the adult. Therefore, we generated the *nubGal4 UASGFP UASgrnd*$^{RNAi}$ recombined flies by crossing *nubGal4 UASGFP* flies with the *UASgrnd*$^{RNAi}$ flies, and crossing putative recombinants with *GMRGal4 UASegr* to score for the rescue of the small eye phenotype to establish recombinant lines (4, 14) .

**Generation FLP-out clones**: The flp-out clones were generated by crossing *y w; Act>y+>Gal4 UASGFP* flies with *yw hsFlp; UASYki*$^{3SA}$*; UASscrib*$^{RNAi}$, and F1 larvae (48 hr after egg laying) were heat shocked at 37˚C for 5 min to induce somatic clones. This heat shock regimen was followed to induce *Yki*$^{3SA}$ *scrib*$^{RNAi}$ clones alone or in combination with other transgenes described in the manuscript. For the *Ras*$^{V12}$*scrib*$^{RNAi}$ flp-out clones, F1 larvae obtained from crossing *y w; Act>y+>Gal4 UASGFP* flies with *yw hsFlp; UASRas*$^{V12}$*; UASscrib* $^{RNAi}$ were heat shocked for 3 min at 37˚C. This shorter heat-shock protocol was used for all 'flp-out' clone experiments involving *Ras*$^{V12}$*scrib*$^{RNAi}$ genetic background. All the crosses were maintained at 25˚C. Wing imaginal discs from wandering third instar larvae were used for analysis.

**Immunohistochemistry:** Third instar larvae were dissected in 1X phosphate buffered saline (PBS) and fixed in 4% paraformaldehyde for 20min. Samples were washed in 1X PBS containing 0.2% Triton X-100 (PBST), blocked for 1hr. with PBST containing 2% Normal Donkey Serum, followed by overnight incubation in primary antibody at 4˚C. Following this, the primary antibody was removed and samples were washed in PBST before incubation in secondary antibody for 2hr at RT. Unbound antibodies were washed and samples mounted in the VectaShield mounting medium (Vector Labs) and scanned using confocal microscopy (Olympus Fluoview 1000, 3000).

**Supplementary Figure information**: The list of genotypes for all panels are listed below. For quantification Two-tailed, unpaired student T-test with $n \geq 5$, 95% confidence was performed using Graphpad Prism 5.

**Figure 1: Expression of Oncogenic Yki (*Yki*$^{3SA}$) in *scrib* mutant cells forms invasive tumors**

(A) *yw hsFLP; nubGal4,UASGFP; TM3/TM6B*,

(A’,B) *yw hsFLP; nubGal4, UASGFP/UASscrib*$^{RNAi}$*; UASYki*$^{3SA}$*/TM6B*,

(C) *yw hsFLP; AyGal4, UASGFP/ UASscrib*$^{RNAi}$*; UASYki*$^{3SA}$*/puc*$^{E69-}$*lacZ*,

(D) *yw hsFLP; AyGal4, UASGFP/ UASRas*$^{V12}$*; UASscrib*$^{RNAi}$*/puc*$^{E69}$*-lacZ*,

(E) *yw hsFLP; AyGal4, UASGFP/ UASscrib*$^{RNAi}$ *; UASYki*$^{3SA}$*/TM6B,*

(F) *yw hsFLP; AyGal4, UASGFP/UASRas*$^{V12}$*; UASscrib*$^{RNAi}$*/TM6B*

**Figure 1: Role of Cact in tumor progression**

(A) *yw hsFLP; AyGal4, UASGFP; TM3/TM6B*,

(B) *yw hsFLP; AyGal4, UASGFP/UASscrib*$^{RNAi}$*; TM3/TM6B*,

(C) *yw hsFLP; AyGal4, UASGFP; UASYki*$^{3SA}$*/TM6B*,

(D) *yw hsFLP; AyGal4, UASGFP/UASscrib*$^{RNAi}$*; UASYki*$^{3SA}$*/TM6B*,

(F,K) *yw hsFLP; AyGal4, UASGFP; UAScact*$^{RNAi}$*/TM6B*,

(G,L) *yw hsFLP; AyGal4, UASGFP/UASscrib*$^{RNAi}$ *; UAScact*$^{RNAi}$*/TM6B*

(H,M) *yw hsFLP; AyGal4, UASGFP; UASYki*$^{3SA}$*/UAScact*$^{RNAi}$,

(I, N) *yw hsFLP; AyGal4, UASGFP/*$UASscrib^{RNAi}$*; *$UASYki^{3SA}/UAScact^{RNAi}$

**Quantification:** P-values for key comparisons are as follows: (E) p = 0.0017 (Wild-type and $Yki^{3SA}$ clones), p = 0.0160 (Wild-type and $Yki^{3SA}scrib^{RNAi}$), (J) p = 0.061 (Wild-type and $Yki^{3SA}scrib^{RNAi}$), p = 0.048 ($Yki^{3SA}scrib^{RNAi}$ and $Yki^{3SA}scrib^{RNAi}cact^{RNAi}$ clones) (O) p = 0.0228 (Wild-type and $Yki^{3SA}scrib^{RNAi}$), p = 0.0292 ($Yki^{3SA}scrib^{RNAi}$ and $Yki^{3SA}scrib^{RNAi}cact^{RNAi}$ clones).

**Figure 2: Effect of downregulating both the TNF receptors on the Tumor progression**

(A, F) *yw hsFLP; AyGal4, UASGFP*; $UASwgn^{RNAi}/TM6B$,

(B, G) *yw hsFLP; AyGal4, UASGFP*/$UASscrib^{RNAi}$*;* $UASwgn^{RNAi}/TM6B$

(C, H) *yw hsFLP; AyGal4, UASGFP;* $UASYki^{3SA}/UASwgn^{RNAi}$,

(D, I) *yw hsFLP; AyGal4, UASGFP*/$UASscrib^{RNAi}$*;* $UASYki^{3SA}/UASwgn^{RNAi}$

**Quantification:** P-values for key comparisons are as follows: (E) p = 0.0061 (Wild-type and $Yki^{3SA}scrib^{RNAi}$ clones), p = 0.0070 (Wild-type and $Yki^{3SA}scrib^{RNAi}wgn^{RNAi}$). (G) p = 0.0228 (Wild-type and $Yki^{3SA}scrib^{RNAi}$ clones), p = 0.0145 (Wild-type and $Yki^{3SA}scrib^{RNAi}wgn^{RNAi}$), (K) p = 0.0010 ($Yki^{3SA}scrib^{RNAi}$ and $Yki^{3SA}scrib^{RNAi}grnd^{RNAi}$), p = 0.0070 ($Yki^{3SA}scrib^{RNAi}$ and $Yki^{3SA}scrib^{RNAi}grnd^{RNAi}wgn^{RNAi}$). (L) No significant difference was found between $Yki^{3SA}scrib^{RNAi}$ and $Yki^{3SA}scrib^{RNAi}grnd^{RNAi}$, $Yki^{3SA}scrib^{RNAi}wgn^{RNAi}$ or $Yki^{3SA}scrib^{RNAi}grnd^{RNAi}wgn^{RNAi}$.

**Figure 4: JNK is the key regulator of tumor invasiveness**

(A, E, H) $UASbsk^{DN}$*/yw hsFLP; AyGal4, UASGFP/*$UASscrib^{RNAi}$*;* $UASYki^{3SA}/TM6B$,

(B, F) $UASbsk^{DN}$*/yw hsFLP; AyGal4, UASGFP/*$UASscrib^{RNAi}$*;* $UASYki^{3SA}/UASwgn^{RNAi}$

**Quantification:** P-values for key comparisons are as follows: (C),p= 0.0005 (wild-type and $Yki^{3SA}scrib^{RNAi}$), p= 0.0243 ($Yki^{3SA}scrib^{RNAi}$ and $Yki^{3SA}scrib^{RNAi}$ $bsk^{DN}$), p= 0.0273 ($Yki^{3SA}scrib^{RNAi}$

and *Yki*$^{3SA}$*scrib*$^{RNAi}$ *bsk*$^{DN}$ *wgn*$^{RNAi}$), (D) p= 0.0015 (*Yki*$^{3SA}$*scrib*$^{RNAi}$ and *Yki*$^{3SA}$*scrib*$^{RNAi}$ *bsk*$^{DN}$), p= 0.0027 (*Yki*$^{3SA}$*scrib*$^{RNAi}$ and *Yki*$^{3SA}$*scrib*$^{RNAi}$ *bsk*$^{DN}$ *wgn*$^{RNAi}$). (G) p= 0.0359 (*Yki*$^{3SA}$*scrib*$^{RNAi}$ and *Yki*$^{3SA}$*scrib*$^{RNAi}$ *bsk*$^{DN}$), p= 0.0351 (*Yki*$^{3SA}$*scrib*$^{RNAi}$ and *Yki*$^{3SA}$*scrib*$^{RNAi}$ *bsk*$^{DN}$ *wgn*$^{RNAi}$). (I) No significant difference between *Yki*$^{3SA}$*scrib*$^{RNAi}$ and *Yki*$^{3SA}$*scrib*$^{RNAi}$ *bsk*$^{DN}$. p= 0.0160 (wild and *Yki*$^{3SA}$*scrib*$^{RNAi}$), p= 0.0020 (*Yki*$^{3SA}$*scrib*$^{RNAi}$ and *Yki*$^{3SA}$*scrib*$^{RNAi}$ *bsk*$^{DN}$).

**Figure 5: Role of Yki in stimulating Cact and promoting tumorigenesis**

(A) *yw hsFLP; AyGal4, UASGFP/UASp35; UASyki*$^{RNAi}$*/ TM6B*,

(B) *yw hsFLP; AyGal4, UASGFP/UASp35; UASyki*$^{RNAi}$*/UASscrib*$^{RNAi}$,

(C) *yw hsFLP; AyGal4, UASGFP/UASp35; UASyki*$^{RNAi}$*/UASRas*$^{V12}$,

(D) *yw hsFLP; AyGal4, UASGFP/UASscrib*$^{RNAi}$*; UASyki*$^{RNAi}$*/UASRas*$^{V12}$,

(H) *yw hsFLP; nubGal4, UASGFP/CyO; UASsd*$^{RNAi}$*/TM6B*,

(I) *yw hsFLP; nubGal4, UASGFP/UASscrib*$^{RNAi}$*; UASsd*$^{RNAi}$*/TM6B*,

(J) *yw hsFLP; nubGal4, UASGFP/CyO; UASsd*$^{RNAi}$*/UASRas*$^{V12}$,

(K) *yw hsFLP; nubGal4, UASGFP/UASscrib*$^{RNAi}$ *; UASsd*$^{RNAi}$*/UASRas*$^{V12}$

**Quantification:** P-values for key comparisons are as follows: (E) p <0.0001 (*Ras*$^{V12}$*scrib*$^{RNAi}$ and *Ras*$^{V12}$*scrib*$^{RNAi}$*yki*$^{RNAi}$) (F) p = 0.0077(Wild-type and *Ras*$^{V12}$*scrib*$^{RNAi}$), p = 0.0081(Wild-type and *Ras*$^{V12}$*scrib*$^{RNAi}$*yki*$^{RNAi}$), p = 0.0017 (*Ras*$^{V12}$*scrib*$^{RNAi}$ and *Ras*$^{V12}$*scrib*$^{RNAi}$*yki*$^{RNAi}$) (G) p = 0.0020 (Wild type and *Ras*$^{V12}$*scrib*$^{RNAi}$), p = not-significant (Wild type and *Ras*$^{V12}$*scrib*$^{RNAi}$*yki*$^{RNAi}$), p = 0.0062 (*Ras*$^{V12}$*scrib*$^{RNAi}$ and *Ras*$^{V12}$*scrib*$^{RNAi}$*yki*$^{RNAi}$) (L) p = 0.0034 (Wild-type and *nub>Ras*$^{V12}$*scrib*$^{RNAi}$), p = non-significant (Wild-type and *nub>Ras*$^{V12}$*scrib*$^{RNAi}$*sd*$^{RNAi}$), p = 0.0032 (*nub>Ras*$^{V12}$*scrib*$^{RNAi}$ and *nub>Ras*$^{V12}$*scrib*$^{RNAi}$*yki*$^{RNAi}$).

**Supplementary Figure Legends:**

**Figure S1 linked to Fig 1: Expression of Oncogenic Yki (*Yki$^{3SA}$*) in *scrib* mutant cells forms invasive tumors.**

(**A**) Overgrowth in *nub>Yki$^{3SA}$scrib$^{RNAi}$* third instar larval wing disc shown at 10X magnification. (**B**) pJNK expression (red, grey) in GFP-labelled clones (green) in *AyGal4>GFP* clones showing jagged edges of the clone. (**C**) Thickness of the clones representing the multi-layered structure of indicated genotypes n = 6, p = 0.0005 (wild-type and *Yki$^{3SA}$scrib$^{RNAi}$*), p = 0.0077(Wild-type and *Ras$^{V12}$scrib$^{RNAi}$*). Panels below show YZ sections used to measure clone size. (**D**) Graph shows quantification of pJNK expression in *Yki$^{3SA}$scrib$^{RNAi}$* tumor (GFP expressing clone) and normal cells (non-GFP expressing cells outside the clone), n =5, p = 0.0032. Panels show expression of (**E**) pJNK (red, grey) in *Yki$^{3SA}$scrib$^{RNAi}$*, and (**F**) MMP1 expression (red, grey) in *AyGal4>GFP* clones (green).

**Figure S2 linked to Fig. 2: Role of Cact in tumor progression.**

Cact expression (red, grey) in GFP-labelled clones (green) from (**A**) *AyGal4> Ras$^{V12}$scrib$^{RNAi}$*, and (**C**) *nub>Yki$^{3SA}$scrib$^{RNAi}$* wing discs. (**B**) Graph shows comparison of Cact expression in wild-type and *Ras$^{V12}$scrib$^{RNAi}$* clones, n=5, p= 0.0020. (**D**) Graph shows quantification of pJNK expression in clones of indicated genotypes**. (E)** Thickness of the clones representing the multi-layered structure of indicated genotypes, n = 6, p = 0.0005 (wild-type and *Yki$^{3SA}$scrib$^{RNAi}$*), p = 0.0003 (Wild-type and *Yki$^{3SA}$scrib$^{RNAi}$cact$^{RNAi}$*). (**F**) Graph shows quantification of MMP1 expression in clones of indicated genotype**. (G)** pJNK expression quantified in indicated genotype to test effects of downregulation of Cact, n=5, p<0.0001 (wild-type and

$nub>Yki^{3SA}scrib^{RNAi}$) p= 0.0028 ($nub>Yki^{3SA}scrib^{RNAi}$ and $nub>Yki^{3SA}scrib^{RNAi}cact^{RNAi}$) (**H**) Graph shows effects of Cact downregulation on MMP1 expression quantified for indicated genotypes, n=5, p= 0.0002 (wild-type and $nub>Yki^{3SA}scrib^{RNAi}$) p= 0.0173 ($nub>Yki^{3SA}scrib^{RNAi}$ and $nub>Yki^{3SA}scrib^{RNAi}cact^{RNAi}$).

**Figure S3 linked to Fig. 3: Quantification of Effect of downregulating both TNF receptors on Tumor progression**

(**A**) Comparison of clone size between $scrib^{RNAi}$, $scrib^{RNAi}cact^{RNAi}$ and $scrib^{RNAi}wgn^{RNAi}$, n=5, p= 0.0195 ($scrib^{RNAi}$ and $scrib^{RNAi}wgn^{RNAi}$) Graph shows quantification of pJNK (**B)** and MMP1 (**C**) expression in clones of indicated genotype**.** (**D**) Graph shows MMP1 expression in $Yki^{3SA}wgn^{RNAi}$ clones, n=5, p= 0.0028 (wild-type and $Yki^{3SA}wgn^{RNAi}$).

**Figure S4 linked to Fig. 3: Effect of downregulating both TNF receptors on Tumor progression.**

Panels show expression of pJNK (red, grey) (**A,C**) and MMP1(red, grey) (**B,D**) in wing discs of indicated genotype at 40X magnification. Downregulation of (**A, B**) *grnd* or (**C, D**) *wgn* in the GFP labelled cells (green) showed decrease in pJNK expression (**A,C**) but not MMP1 expression (**B,D**).

**Figure S5 linked to Fig. 5: Role of Yki in stimulating Cact and promoting tumorigenesis.**

Panels show pJNK (red, grey) (**A,C**) and MMP (red, grey) (**B,D**)expression in GFP labelled (**A,B**) $Ras^{V12}scrib^{RNAi}yki^{RNAi}$ clones, or (**C,D**) in $Ras^{V12}scrib^{RNAi}sd^{RNAi}$ wing discs.

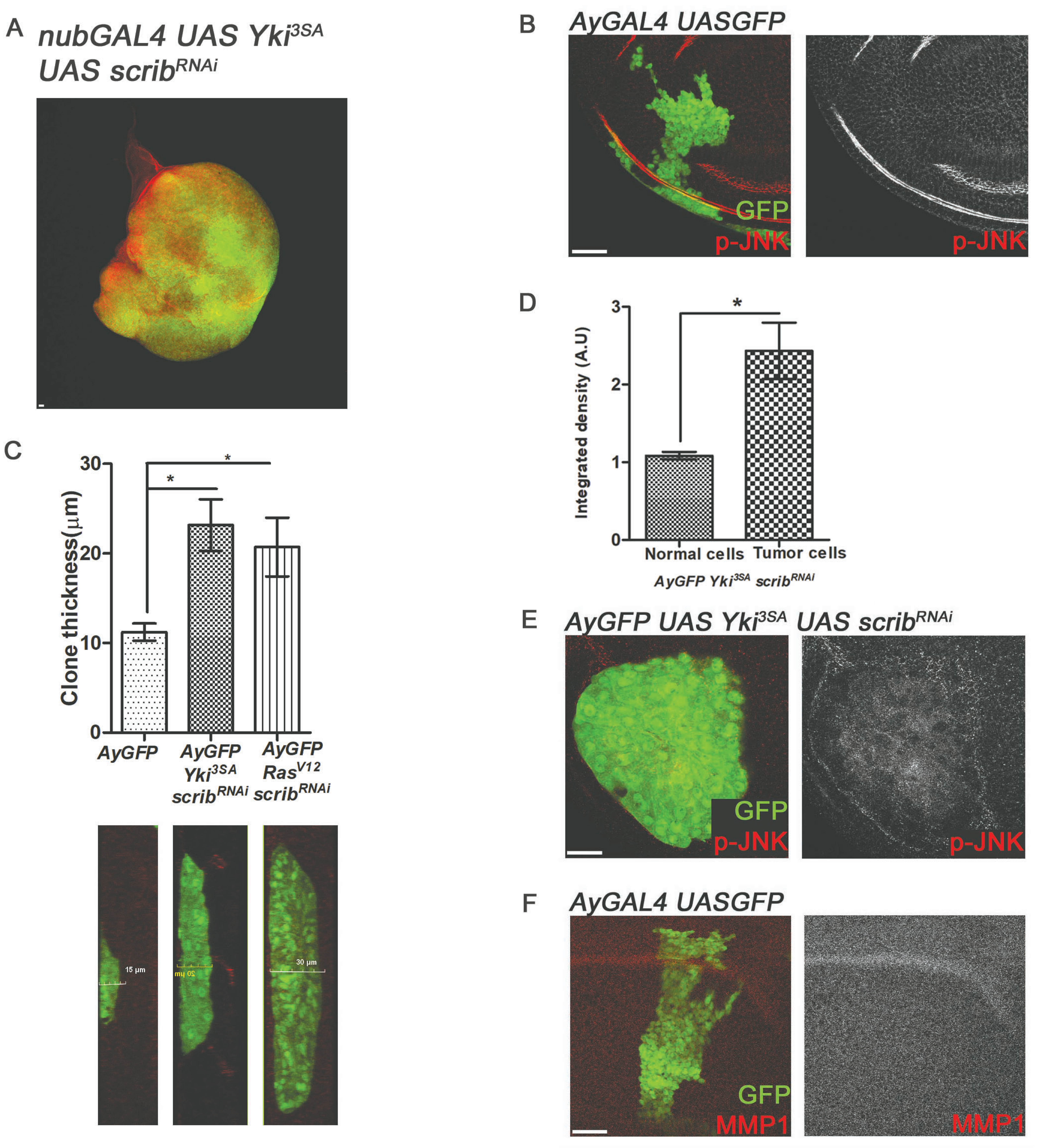


Snigdha et al Figure S1

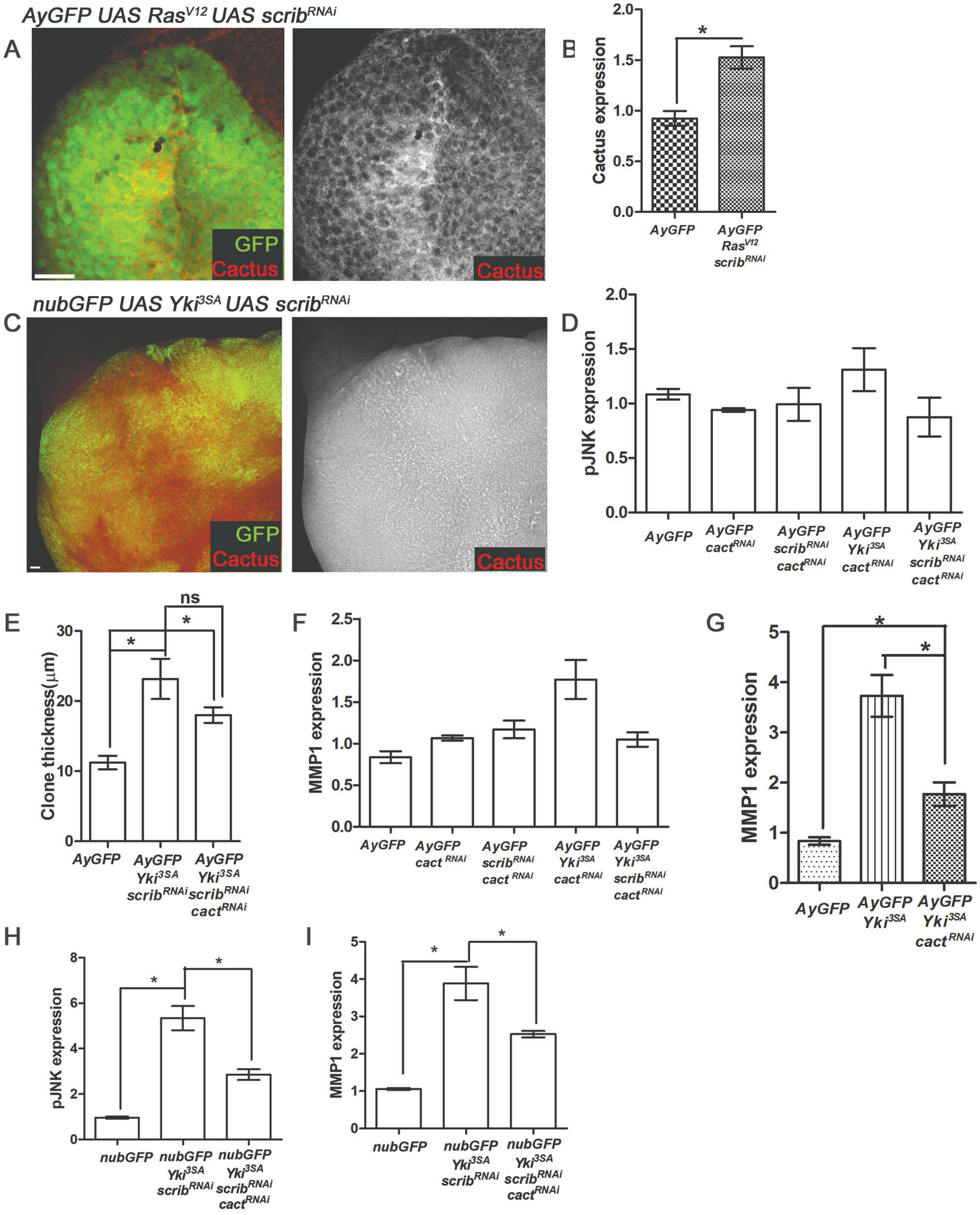


Snigdha et al Figure S2

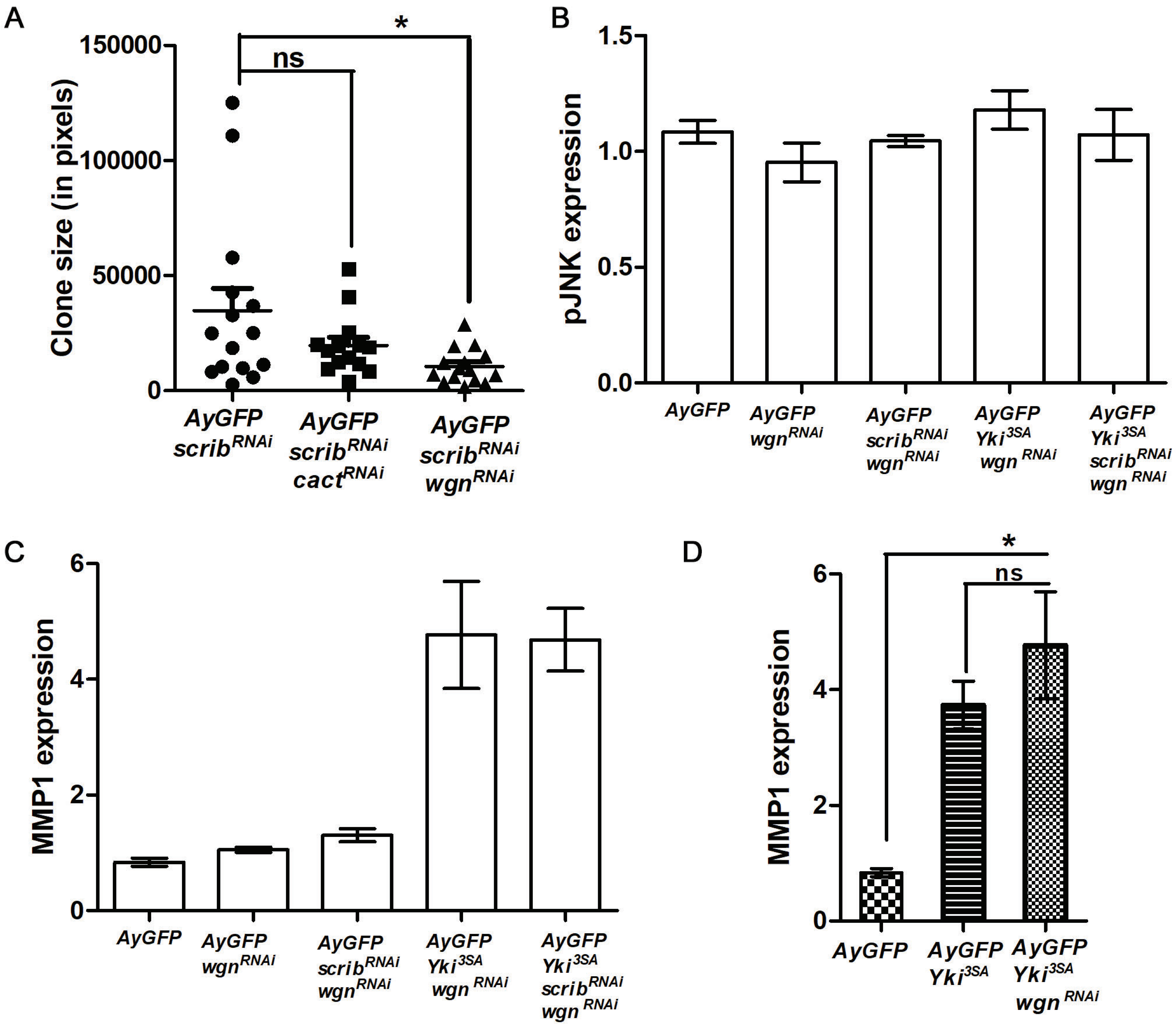


Snigdha et al Figure S3

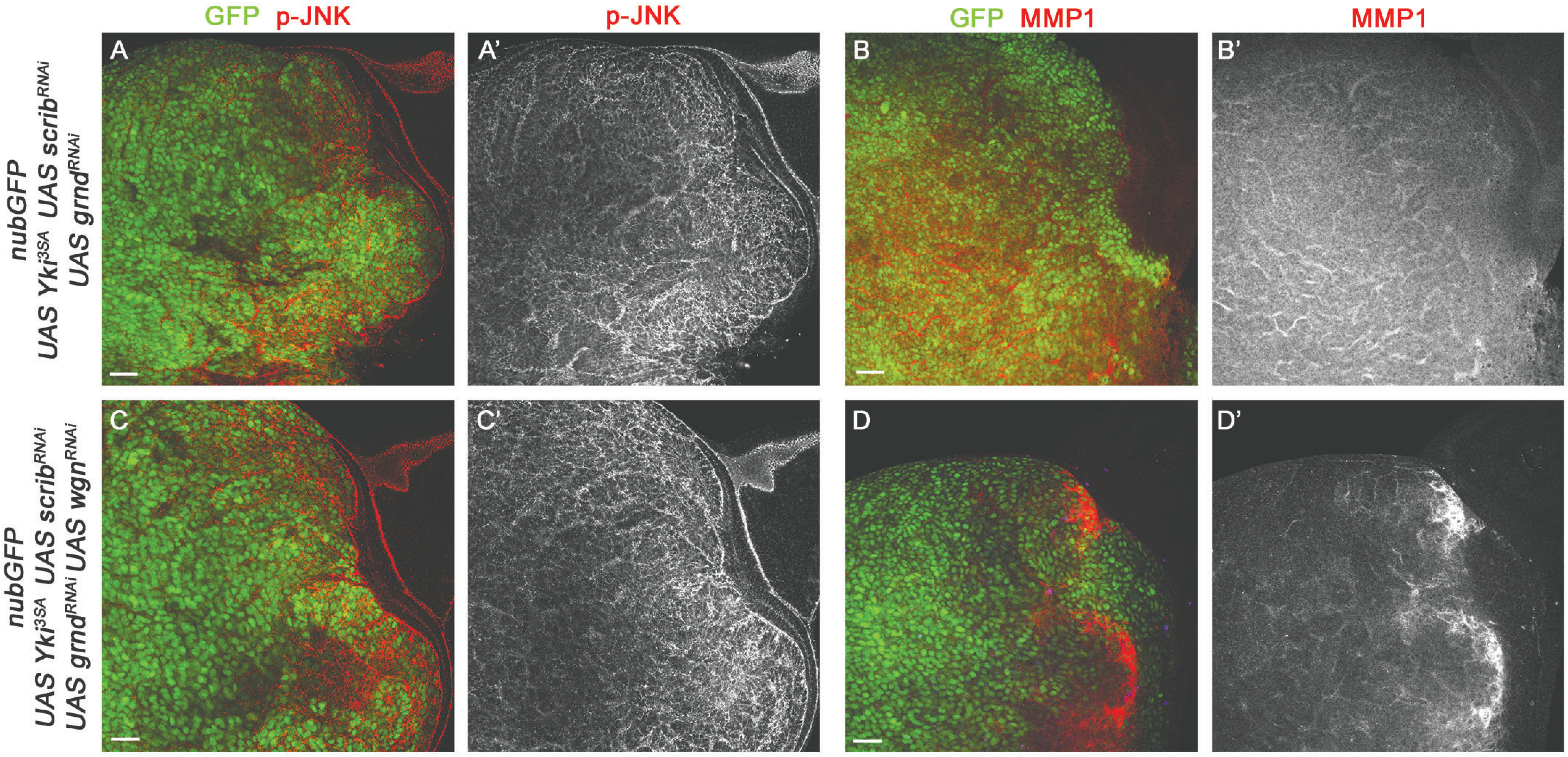


Snigdha et al Figure S4

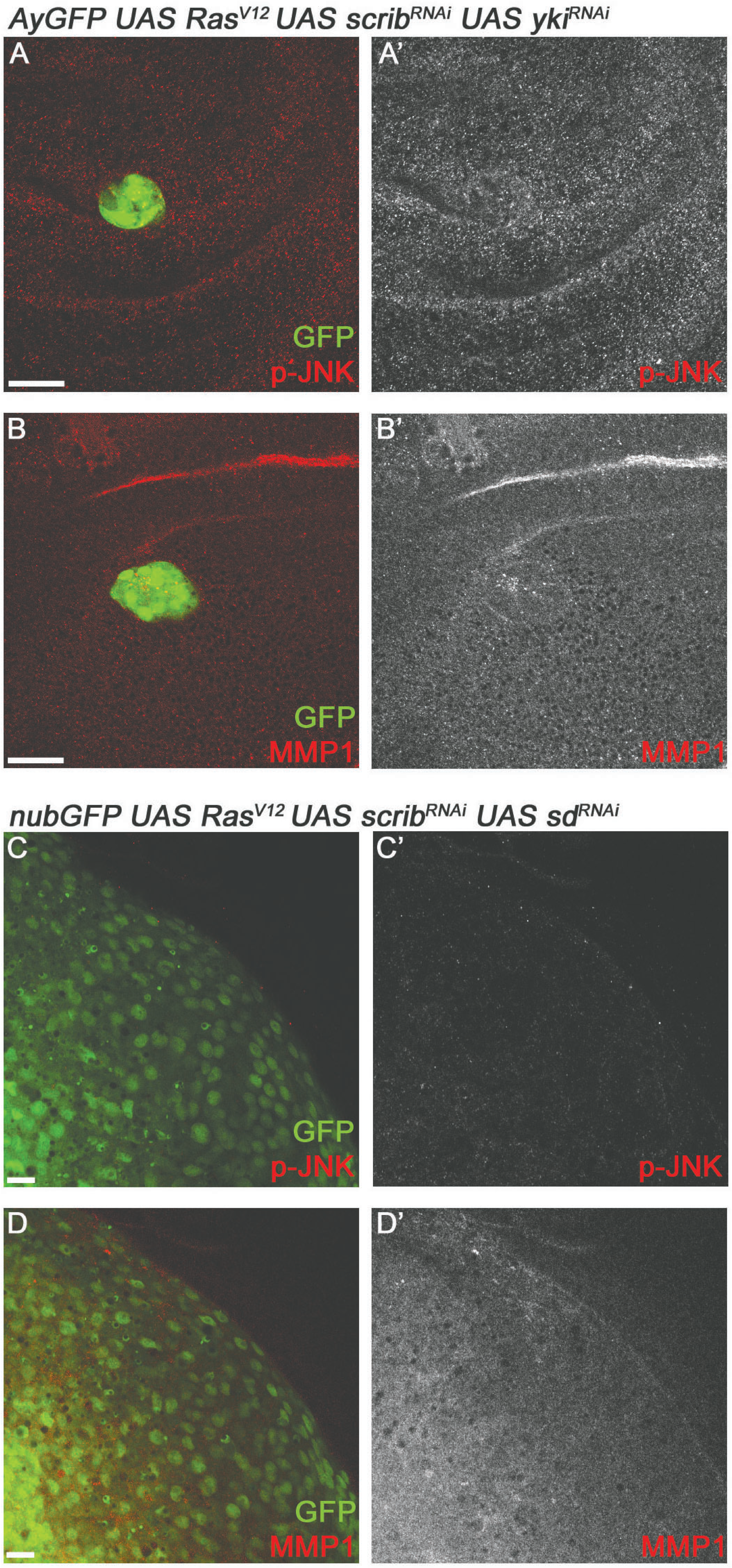


Snigdha et al Figure S5
